# Structural basis for the transport and regulation mechanism of the Multidrug resistance-associated protein 2

**DOI:** 10.1101/2024.06.24.600277

**Authors:** Eriko Koide, Harlan L. Pietz, Jean Beltran, Jue Chen

## Abstract

Multidrug resistance-associated protein 2 (MRP2) is an ATP-powered exporter important for maintaining liver homeostasis and a potential contributor to chemotherapeutic resistance. Deficiencies in MRP2 function are associated with Dubin-Johnson Syndrome and increased vulnerability to liver injury from cytotoxic drugs. Using cryogenic electron microscopy (cryo-EM), we determined the structures of human MRP2 in three conformational states: an autoinhibited state, a substrate-bound pre-translocation state, and an ATP-bound post-translocation state. These structures show that MRP2 functions through the classic alternating access model, driven by ATP binding and hydrolysis. Its cytosolic regulatory (R) domain serves as a selectivity gauge, wherein only sufficiently high concentrations of substrates can effectively compete with and disengage the R domain to initiate transport. Comparative structural analyses of MRP2 in complex with different substrates reveal how the transporter recognizes a diverse array of compounds, highlighting the transporter’s role in multidrug resistance.

## Introduction

Multidrug resistance-associated protein 2 (MRP2) is an ATP-Binding Cassette (ABC) transporter expressed in the liver, intestines, kidney, and placenta^1,2^. Patients deficient in MRP2 function develop Dubin-Johnson Syndrome, a disease caused by an excess of the bile pigment bilirubin in the liver cells^3–5^. Mutations that decrease the amount of functional MRP2 have also been associated with greater incidence of toxic liver injury during administration of cytotoxic drugs^6^. MRP2 knockout mice exhibit hyperbilirubinemia, reduced bile flow, reduced biliary glutathione excretion, and an increase in liver size^7^. These data indicate that MRP2 plays a role in maintaining liver homeostasis by excreting potentially toxic molecules. In addition to endogenous substrates such as bile components and conjugated metabolites, MRP2 also recognizes and transports xenobiotic molecules including anthracyclines and vinca alkaloids^1^. MRP2 upregulation is associated with poor prognosis in esophageal squamous cell carcinoma, which likely arises from the ability of MRP2 to transport many chemotherapeutic molecules^8^. Hence, MRP2 poses a persistent challenge in the pharmacology of multidrug resistance-associated proteins: while inhibition holds therapeutic promise to overcome drug resistance in chemotherapy, it also carries the risk of inducing toxicity.

MRP2 is a single polypeptide consisting of an N-terminal transmembrane domain (TMD0) and a transporter core composed of two transmembrane domains (TMD1 and TMD2) along with two nucleotide-binding domains (NBD1 and NBD2) (Figure 1A). The function of TMD0 in MRP2 remains unclear and is only conserved in a small number of ABC transporters. Studies deleting TMD0 in MRP1, a homolog of MRP2, showed no functional effects on ATP hydrolysis and substrate transport^9,10^. The transporter core, on the other hand, is highly conserved among ABC exporters. In these exporters, the two TMDs form a substrate translocation pathway, while the NBDs bind and hydrolyze ATP^11^. Additionally, MRP2 contains a 100-residue long linker between NBD1 and TMD2, resembling the regulatory (R) domain of the cystic fibrosis transmembrane regulator (CFTR).

**Figure 1.**
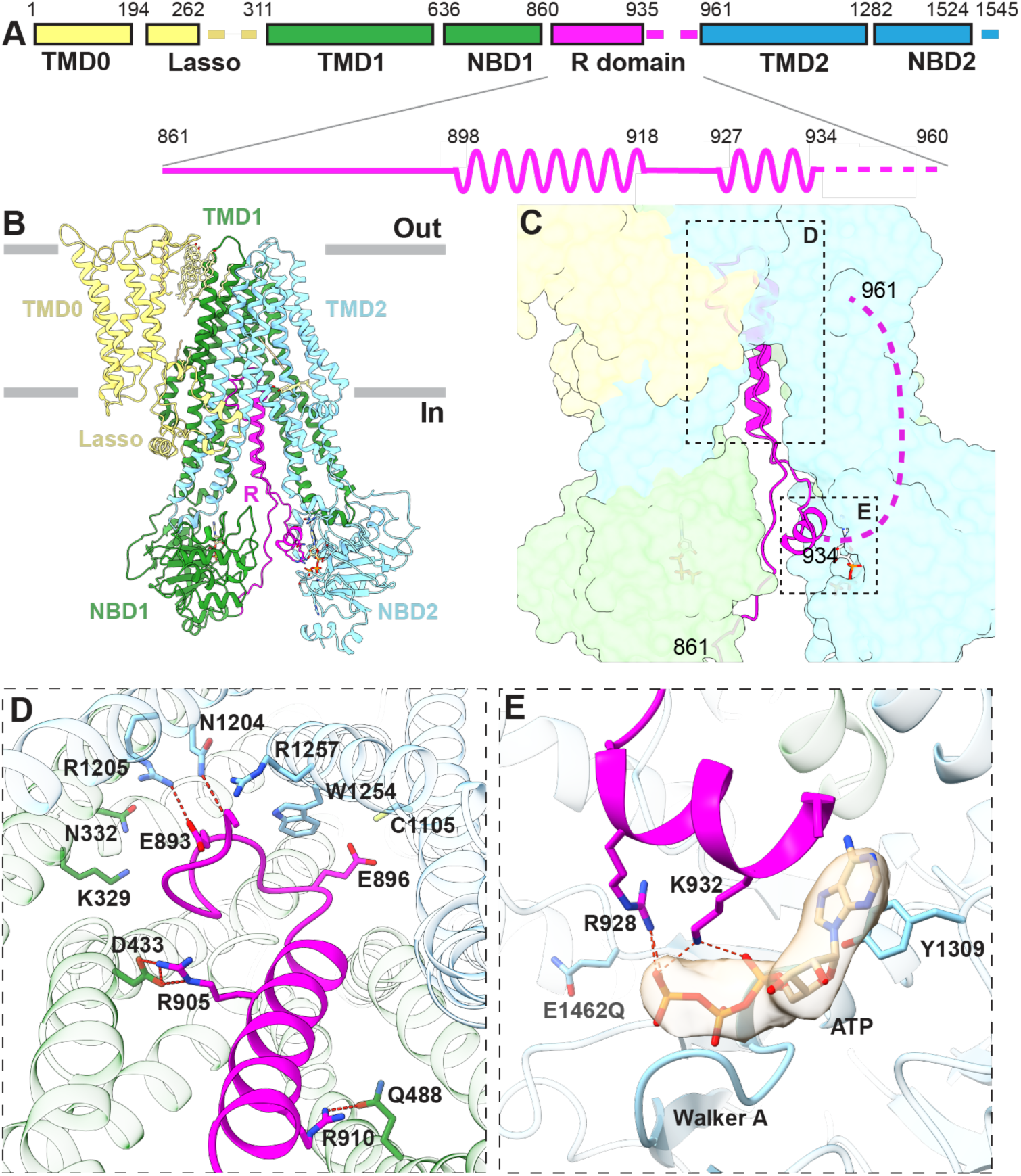
The resting state of MRP2 reveals an auto-inhibitory R domain. (A) Domain organization of MRP2 with the secondary structure of the R domain depicted. (B) Ribbon representation of the structure of MRP2(E1462Q) in the inward-facing, ATP-bound conformation. (C) Zoomed-in view with TMDs and NBDs represented as surfaces and the R domain in ribbon. Boxes indicate the positions of the two interfaces illustrated in panels D and E. (D) R domain inserts into the substrate-binding site as viewed from the cytoplasm. Hydrogen bonds are indicated by red dashed lines. (E) An ATP molecule binds between the R domain and NBD2. Cryo-EM density of ATP is depicted as a transparent surface. Dashed lines indicate hydrogen bonds.

A fundamental puzzle regarding multidrug resistance is how a single transporter can recognize many unrelated drugs. The mechanisms of other multidrug transporters, including P-glycoprotein (P-gp, ABCB1), ABCG2, and MRP1 have been well-studied. For example, P-gp and ABCG2 both contain a central hydrophobic cavity that many small-molecules, including substrates and inhibitors, can bind largely through van der Waals interactions^12–17^. The substrate-binding site in MRP1 is very different from those in P-gp and ABCG2. It can be divided into two parts: a positively charged region containing many polar functional groups (which was termed the P-pocket) and a largely hydrophobic area capable of interacting with different hydrophobic moieties (the H-pocket)^9^. In addition, many residues in the binding site exhibit side-chain plasticity that would allow for adaptability to a variety of substrates. The mechanism by which MRP2 can recognize many substrates with different chemical structures remains to be elucidated.

In this study, we used cryogenic electron microscopy (cryo-EM) to investigate the structural dynamics of MRP2 throughout its transport cycle. Through structural analysis of MRP2 in auto-inhibited, pre-translocation, and post-translocation states, we observed large scale conformational changes enabling ATP-dependent substrate translocation. Furthermore, by comparing the structures of MRP2 in complex with different substrates, we begin to understand how a single transporter recognizes a diverse array of substrates.

## Results

### The resting state of MRP2 is an auto-inhibited state

We first determined the structure of the wild type (WT) human MRP2 in the absence of substrate and ATP (apo) and observed an inward-facing, NBD-separated conformation (Fig. S1, S3A). We also analyzed the structure of a hydrolysis-deficient variant (E1462Q) in the presence of ATP, which exhibited two conformations with approximately equal populations (Fig. S2): an inward-facing, NBD-separated conformation and an NBD-dimerized conformation which we will discuss later. Despite the different sample preparation conditions, the inward-facing conformations of the WT and the E1462Q variant were very similar: when superimposed, the overall root mean square deviation (RMSD) is 3.1 Å for 1422 Cα positions (Fig. S3B, S3C). Because the structure of the E1462Q variant reveals more complete density of the R domain, we will discuss the inward-facing conformation based on this structure (Fig. 1).

In this conformation, the structures of TMD0 and the transporter core are both well defined. While the five TM helices of TMD0 constitute a compact domain, the twelve TM helices of TMD1 and TMD2 form two domain-swapped, pseudo symmetric bundles, each with an NBD attached in the cytosol (Fig. 1B, S3C). Interactions between TMD0 and the transporter core occur mostly in the extracellular loops and with the lasso motif (Fig. 1B, S3D). In the transmembrane region, TMD0 and the transporter core have few direct contacts; instead, they are bridged by cholesterol and lipid molecules.

A unique feature of MRP2, compared to other MRP molecules, is its structured R domain—the 100-residue linker between NBD1 and TMD2 (Fig. 1B, C, Fig. S2C). The N-proximal half of the R domain (residues 861-893) forms a long loop along the inner surface of NBD1, extending nearly 90 Å into the TM cavity (Fig. S3E). Residues 894-897 form a sharp turn at the top of the TM cavity, and residues 898-918 form an α-helix between the two TM bundles (Fig. 1C-D). The remainder of the R domain extends into the cytosol and forms a short helical segment at the inner surface of NBD2 (Fig. 1E). Density for the C-terminal 25 residues of the R domain was absent, indicating the flexibility of the linker connecting to TMD2.

Two interfaces between the R domain and the transporter core may have important mechanistic implications. Near the apex of the TM cavity, the loop-turn-helix motif of the R domain engages in hydrogen bonding and extensive van der Waals interactions with TMD1 and TMD2 (Fig. 1D). Many residues at this interface, including R1205, N1204, R1257, W1254, and N332, are involved in substrate binding (discussed later, in Fig. 2). This configuration is reminiscent of the inhibited structure of the peptide transporter TAP, in which a viral inhibitor ICP47 forms a helical hairpin inserting into the intracellular openings of the transporter to preclude substrate binding and conformational change necessary for transport function^18,19^. It seems likely that, in the absence of substrate, MRP2 rests in an auto-inhibited state to minimize futile ATP hydrolysis.

**Figure 2.**
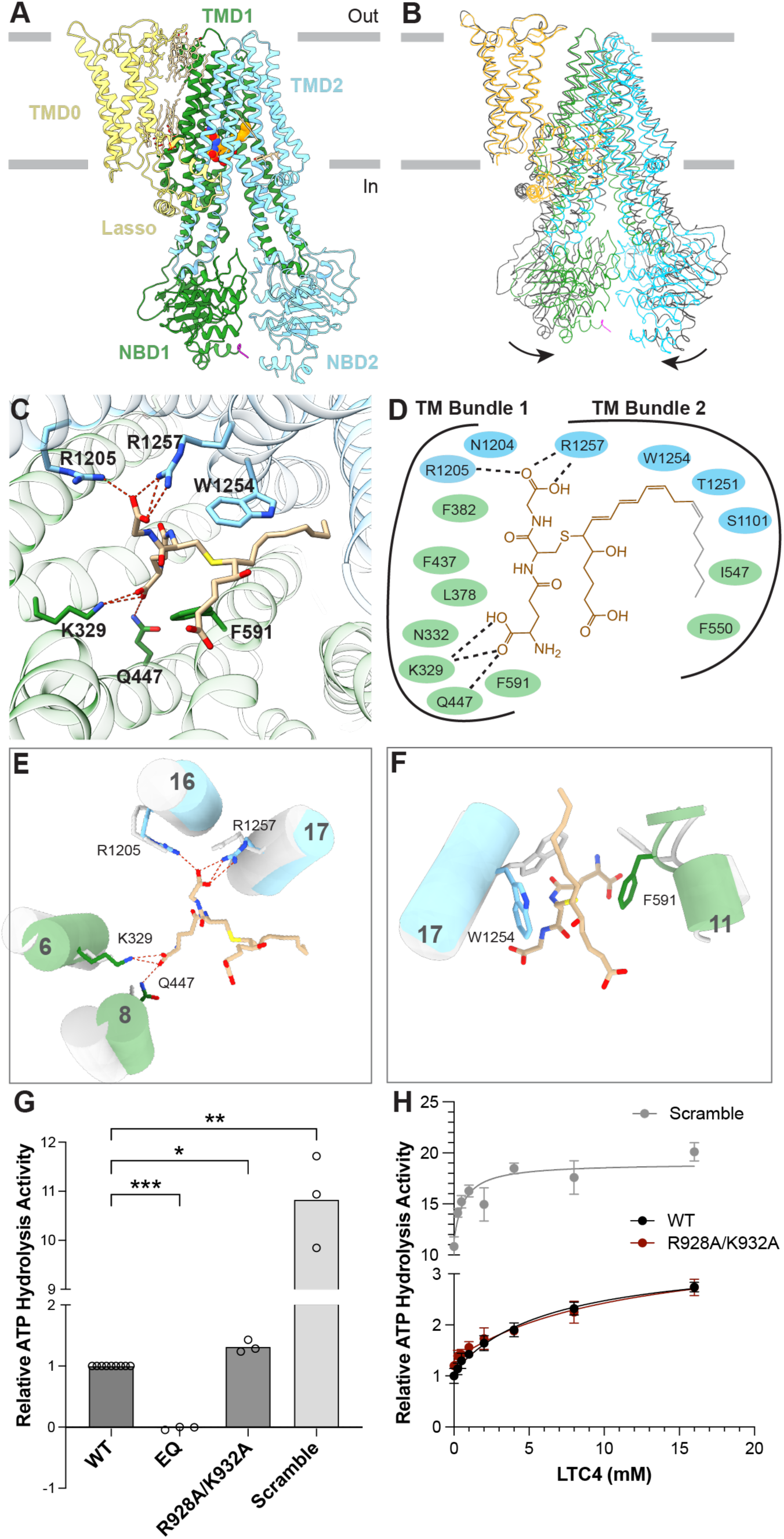
LTC4 binding releases R domain inhibition. (A)The overall structure of the LTC4-bound MRP2. (B) Superposition of the LTC4-bound structure (colored) with that of the substrate-free MRP2 (gray). Arrows indicate the inward movement upon LTC4 binding. (C) Molecular details of LTC4 binding, viewed from the cytosol. Hydrogen bonds between LTC4 and MRP2 are indicated by dashed lines. (D) Schematic drawing of the LTC4-MRP2 interactions. The unresolved hydrophobic tail of LTC4 is shown in gray. Hydrogen bonds are depicted as dashed lines. (E) (F) Local conformational changes at the LTC4-binding site, colored as in panel C and overlayed with inward-facing, substrate-free MRP2 (gray). (G) Relative ATPase activity of MRP2 variants. WT: wild type MRP2, EQ: the E1462Q variant, R928A/K932A: double mutation; and Scramble: a variant in which the R domain sequence is randomly scrambled. ATP hydrolysis rates were measured in the presence of 4 mM ATP/magnesium at a protein concentration of 75 nM, except for the E1462Q variant, which was 500 nM. Bar height represents the mean value of three separate measurements (unfilled dots). Statistical significance was calculated using matched mixed-effects analysis on the log-transformed relative ATPase activity values. The *p* value of MRP2(E1462Q) is 0.0003, for R928A/K932A is 0.0432, and for Scramble is 0.0042 (*P<0.05, **P<0.01, ***P<5×10^−4^). (H) ATP hydrolysis activity of MRP2 variants, normalized to the basal activity of wild type MRP2 at saturating ATP concentration. Each datapoint represents the mean value of 3 replicates. Error bars represent standard deviation. The curve was fitted with the quadratic binding formula for substrate binding (see methods). The Km values for LTC4: WT: 6.3 uM; R928A/K932A: 11.2 uM; and Scramble: 0.6 uM.

The other interface occurs at the NBD2 surface, where an ATP molecule is coordinated by the Walker A and B motifs of NBD2, along with the R-domain residues R928 and K932 (Fig. 1E). This interface is only observed in the structure of the E1462Q variant, in which 5 mM ATP was included in the sample. In NBD1, clear densities for ATP and Mg were also observed, but the R-domain is not involved in ATP binding (Fig. S3F). As the intracellular ATP concentration is estimated to be 1-10 mM, we suggest that under physiological conditions, ATP is likely bound to both NBDs, further stabilizing the R domain in the auto-inhibited conformation.

### Substrate-binding alleviates auto-inhibition

To understand how substrate is recognized by MRP2 and the effect of substrate binding on the autoinhibitory R domain, we determined the cryo-EM structure of WT MRP2 in the presence of leukotriene C4 (LTC_4_), a pro-inflammatory signaling molecule that is transported by multiple MRP transporters (Fig. 2, Fig. S4A-C). Cryo-EM reconstruction obtained in the presence of 40 μM LTC4 revealed the same structure as the auto-inhibited state; only in the presence of 285 μM LTC4 were we able to observe clear density for LTC_4_ in the substrate-binding site (Fig. S4D). The cryo-EM map, refined to an overall resolution of 2.75 Å, shows no density for the R domain, indicating that the R domain is flexible in this conformation. These results suggest that only at sufficiently high concentrations can the substrate displace the R domain to be transported.

LTC4 binds at the interface of the two TM bundles, triggering a global conformational change that brings the two halves of the transporter core closer together (Fig. 2B, S5A). The inward-swing of the NBDs originates at the substrate-binding site, where TM helix 6, 8, 16, and 17 move towards each other and the side chains of F591 and W1254 reposition to contact LTC_4_ (Fig. 2C,D). Previous studies of MRP1 have shown that LTC4 induces a similar conformational change and accelerates the transition of MRP1 from an NBD-separated, inward-facing conformation to an NBD-dimerized, outward-facing conformation^20,21^. Given that LTC4 also stimulates the ATPase activity of MRP2 (Fig. 2H), it is plausible that MRP2 operates through a comparable mechanism.

Although LTC_4_ interacts with both MRP1 and MRP2, its affinity for MRP2 is approximately ten times lower than that for MRP1^22^. This difference can be attributed to structural differences at their respective binding sites (Fig. S5B). In MRP2, LTC_4_ is largely stabilized by hydrogen bonds between the GSH moiety and residues K329, Q447, R1205, and R1257 (Fig. 2E, F). Despite the high resolution of the cryo-EM map, density for the lipid tail is mostly absent, suggesting that it is not engaged in specific interactions with MRP2 (Fig. S4D). In comparison, in MRP1, the GSH moiety of LTC4 forms an additional hydrogen bond with Y440. The significance of this interaction is underscored by the Y440F variant of MRP1, which leads to reduced ATPase activity upon LTC4- and GSH-stimulation^23^. In MRP2, the corresponding residue to Y440 is, in fact, a phenylalanine (F437), mirroring the Y440F variant of MRP1. In addition, the hydrophobic tail of LTC4 in MRP1 is stabilized by π-orbital stacking interactions with the side chains of M1092, Y1242, and W553, which are not conserved in MRP2 (Fig. S5B). This difference further contributes to the relative lower affinity of MRP2.

To functionally assess the interplay of substrates and the R domain, we generated two variants of the R domain to test its functional relevance (Fig. 2G). Compared to the WT protein, the R928A/K932A double mutant, which disrupts R-domain coordination of ATP but retains the binding-site interactions, caused a modest yet reproducible increase in basal ATPase activity. Disruption of the interactions with the substrate-binding site led to a more pronounced change. Replacement of the entire R domain (residues 861-960) with a scrambled amino acid sequence (“scramble”) increased the basal ATPase rate by more than 10-fold (Fig. 2G). These results suggest that by disrupting the interactions between the R domain and the transporter core, the R domain is more readily released from its auto-inhibitory role, leading to an elevation in the ATPase activity even in the absence of substrate.

LTC_4_ stimulated the ATP hydrolysis rate of both WT and the two MRP2 variants, but with different potencies (Fig. 2H). The apparent K_m_ of the WT protein is approximately 6.3 uM. While perturbation of the ATP-binding residues (the R928A/K932A variant) had only a minor effect on the K_m_ value, disrupting the interaction at the substrate-binding site (the scramble variant) increased the apparent affinity for LTC_4_ by 10-fold. These data suggest that the R domain functions as an affinity/concentration gauge, selecting substrates with either high affinity or those that accumulate to high intracellular concentrations.

### NBD dimerization enables substrate release to extracellular space

The structure of the NBD-dimerized conformation, determined from the dataset collected for the E1462Q variant in the presence of ATP, provides a structural understanding of how substrate is transported and released to the extracellular space. The map is refined to an overall resolution of 3.4 Å (Fig. S2A,D,E) and shows two copies of ATP and magnesium ion at the NBD dimer interface (Fig. S6A,B). Similar to the inward-facing structures, lipid molecules fill the gap to bridge TMD0 and the transporter core (Fig. S6C). Density for the transmembrane helices near the extracellular leaflet is less well defined, suggesting that these helices are relatively flexible (Fig. S2E, compared to S2C, S1C).

In this conformation, much of the R domain remains unresolved, except for the C-proximal residues 945-960. These residues dock along the intracellular region of the TM helices, stabilized by a network of van der Waals interactions and hydrogen bonds (Fig. 3A,B). In particular, K953, Q955, and G960 of the R domain form hydrogen bonds with R1079 and R1083 of TM helix 14 and with residue R1150 of TM helix 15. A similar configuration was previously observed in MRP1^20^ despite the limited sequence homology between the MRP2 R domain and the MRP1 L1 linker (Fig. S6D).This structural similarity suggests a common role for the linker region in stabilizing the NBD-dimerized conformation.

**Figure 3.**
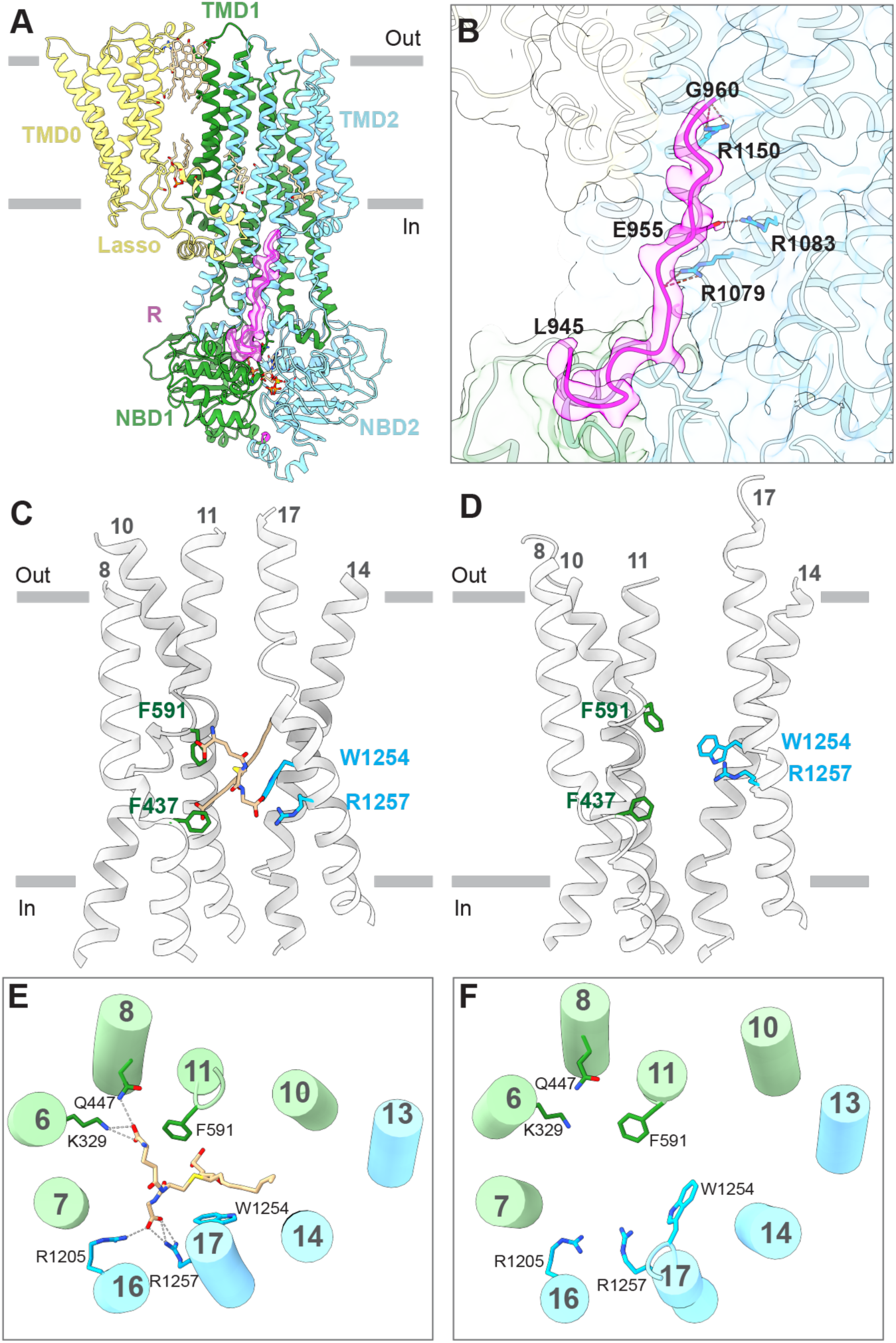
The NBD-dimerized, outward-facing conformation. (A)Ribbon representation of the overall structure. Also shown is the cryo-EM density corresponding to the R domain (magenta surface). (B) Zoomed-in view of the R domain. (C) (D) conformational changes at the substrate-binding site in the inward-facing, LTC4-bound state (C) versus the outward-facing conformation (D). (E) The structure of the substrate-binding site in LTC4-bound conformation, viewed from the extracellular space. (F) The structure of the substrate-binding site in outward-facing conformation, viewed from the extracellular space. Hydrogen bonds are represented by dashed lines.

The TMDs of MRP2 form an outward-facing conformation in which the substrate-binding site is open to the extracellular space and closed off to the cytosol, creating a clear translocation path through which the substrate can be released to the extracellular space (Fig. 3C-F). Several key changes at the substrate-binding site further facilitate substrate release. Upon NBD-dimerization, TM helices 6, 8, and 16 move away from each other, pulling apart residues that interacts with LTC4. Furthermore, R1257 forms a cation-π interaction with W1254, stabilizing W1254 in a conformation that would sterically clash with the substrate (Fig. S6E). These conformational changes have a net effect of reducing the affinity for LTC4 thereby facilitating its release to the extracellular space.

## Discussion

In this manuscript, we present three structures of human MRP2 to elucidate the structural changes associated with its transport cycle and identify a novel autoinhibited conformation. During the preparation of this manuscript, two similar studies were published, with one describing the structure of human MRP2 and the other describing those of rat Mrp2 (rMrp2)^24,25^. Comparing the results obtained in these three independent studies offers a deeper understanding of the MRP2 transport mechanism.

All three studies report a similarly inward-facing, apo conformation with the R domain inserted into the transmembrane substrate-binding site. In addition, we observed the C-proximate region of the R domain that, in the presence of ATP, reaches onto the NBD2 surface to form a composite ATP-binding site (Fig. 1). Perturbation of the R domain binding either by point-mutation or sequence randomization increased the activity of the transporter (^24^ and this study) suggesting that the R domain functions as an auto-inhibitor to down regulate ATP hydrolysis. Only at sufficiently high concentrations can substrate effectively compete for the binding site, promoting the disengagement of the R domain from the inhibitory position. Furthermore, Beis and colleagues have reported higher activities of the fully phosphorylated rMrp2 compared to dephosphorylated or partially phosphorylated protein, suggesting that phosphorylation may regulate the transport activity in a mechanism akin to CFTR^25^. Whether and how phosphorylation of the R domain regulates MRP2, and which kinases are involved, will be the subjects of future studies.

Comparison of the NBD-dimerized, ATP-bound MRP2 structure determined in this study with that of ATP/ADP-bound MRP2 described by Chen and colleagues^24^ reveals that the two structures are nearly identical, with a RMSD of 1.5 Å between 1385 Cα pairs (Fig. S6F). These two structures represent two conformational states: the pre-hydrolysis state with ATP molecules bound at both ATPase sites, and the post-hydrolysis state in which ATP in the catalytically competent site has already been hydrolyzed. The nearly identical structures of these two conformations indicates that substrate is released before ATP hydrolysis, and dissociation of the NBD dimer likely constitutes the rate-limiting step in the transport cycle.

Collectively, the three studies support a mechanism of how ATP hydrolysis is coupled to substrate export (Fig. 4). In the absence of substrate, MRP2 rests in an inward-facing conformation with the R domain inserted between the separated NBDs. At physiological ATP concentrations, ATP is likely bound at the interface of NBD2 and the R domain, further stabilizing the R domain in the auto-inhibitory position (Fig. 4a). When the intracellular concentration of the substrate becomes sufficiently high, it can displace the R domain from the transmembrane binding site (Fig. 4B) to permit NBD dimerization (Fig. 4C). The conformational changes in NBDs are transmitted to the TMDs, opening the translocation pathway to the extracellular space and lowering its affinity so that the substrate will be released (Fig. 4C). ATP hydrolysis followed by ADP release then resets the transporter to its inward-facing configuration, ready again to bind substrate from the cytoplasm.

**Figure 4.**
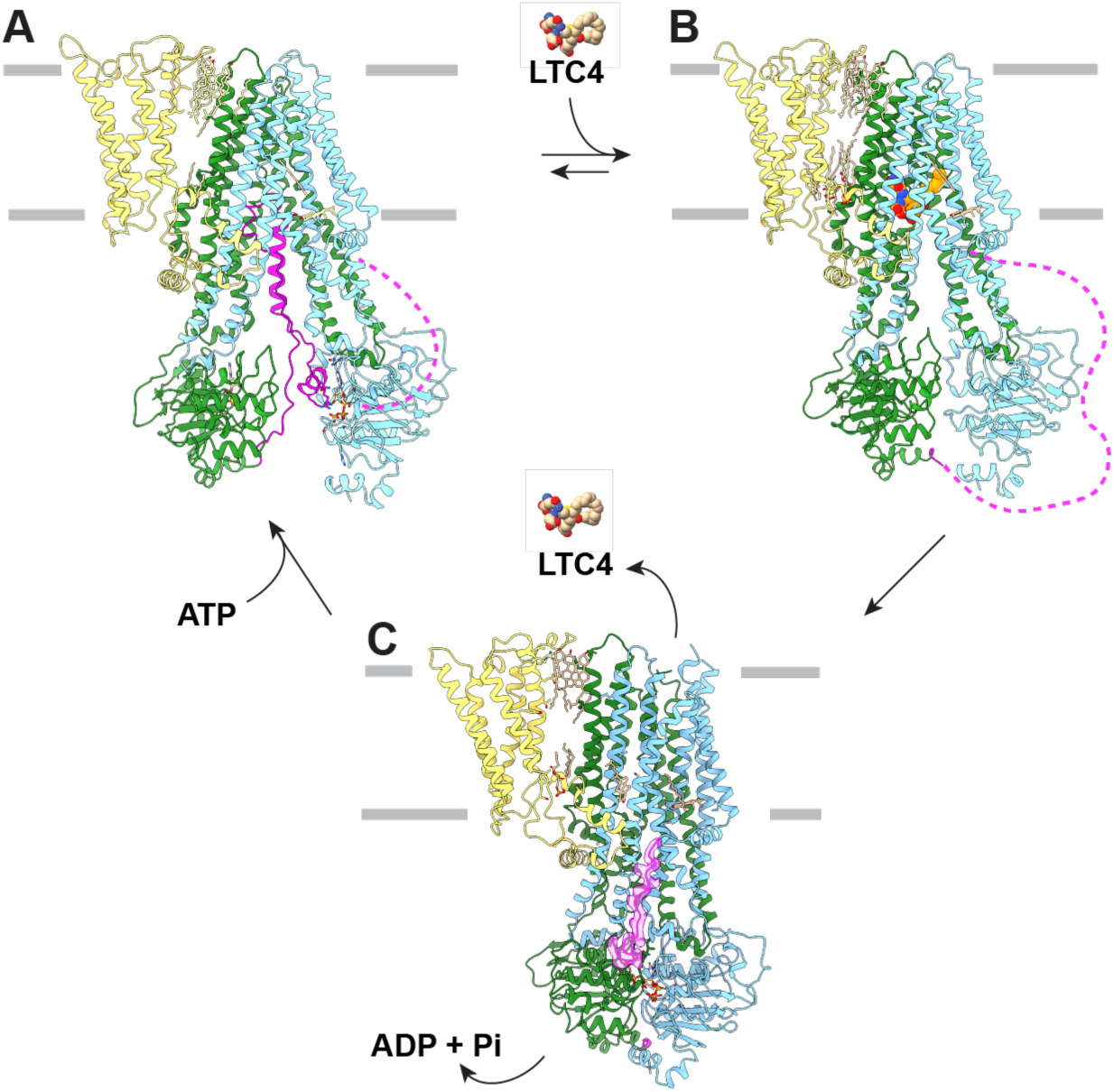
The MRP2 transport cycle. (A) In the absence of substrates, MRP2 rests in an auto-inhibited state. Dotted magenta lines represent unresolved regions of R domain. In this state, ATP-binding does not induce NBD dimerization, but rather stabilizes R domain in the auto-inhibiting position. (B) Substrate displaces R domain from the auto-inhibited conformation. (C) Substrate is released in the NBD-dimerized conformation.

Comparing the three ligand-bound structures of MRP2 (^24,25^ and this study) also provides an opportunity to understand how a single transporter recognizes a spectrum of compounds (Fig. 5). The molecular masses of these three compounds differ by a factor of three, ranging from 285 g/mol (probenecid) to 840 g/mol (BDT). Their chemical structures are also very different: LTC4 is an arachidonic acid derivative conjugated to the tripeptide GSH, BDT is bilirubin conjugated with two taurine moieties, and probenecid is a small sulfonamide. A common feature of these compounds is their bipartite nature, with one portion negatively charged and the other hydrophobic. All three compounds bind to the same pocket in MRP2, which contains a positively charged P-pocket and a mostly hydrophobic H-pocket. The negative moieties of each compound form hydrogen bonds with the same set of residues in the P-pocket (Fig. 5). These hydrogen bonds are critical for ligand recognition, as alanine substitution of a single positively charged residue nearly abolished ATPase stimulation by BDT^24^. The hydrophobic moiety of each compound is encompassed in the H-pocket through van der Waals interactions, and in the cases of BDT and probenecid, a cholesterol molecule was observed “filling in” the space in the H-pocket. Unlike LTC4 or BDT, Probenecid, a drug used to treat gout and gouty arthritis, is inadvertently exported by MRP2^26^. Despite being much smaller than typical MRP2 substrates, two copies of Probenecid occupy the binding site to facilitate its efflux by MRP2^25^. This observation underscores the promiscuous nature of these multidrug transporters and the challenge of avoiding drug resistance conferred by them. Identifying the molecular determinants of ligand recognition and unique characteristics of each transporter will facilitate future drug development efforts aimed at overcoming these challenges.

**Figure 5.**
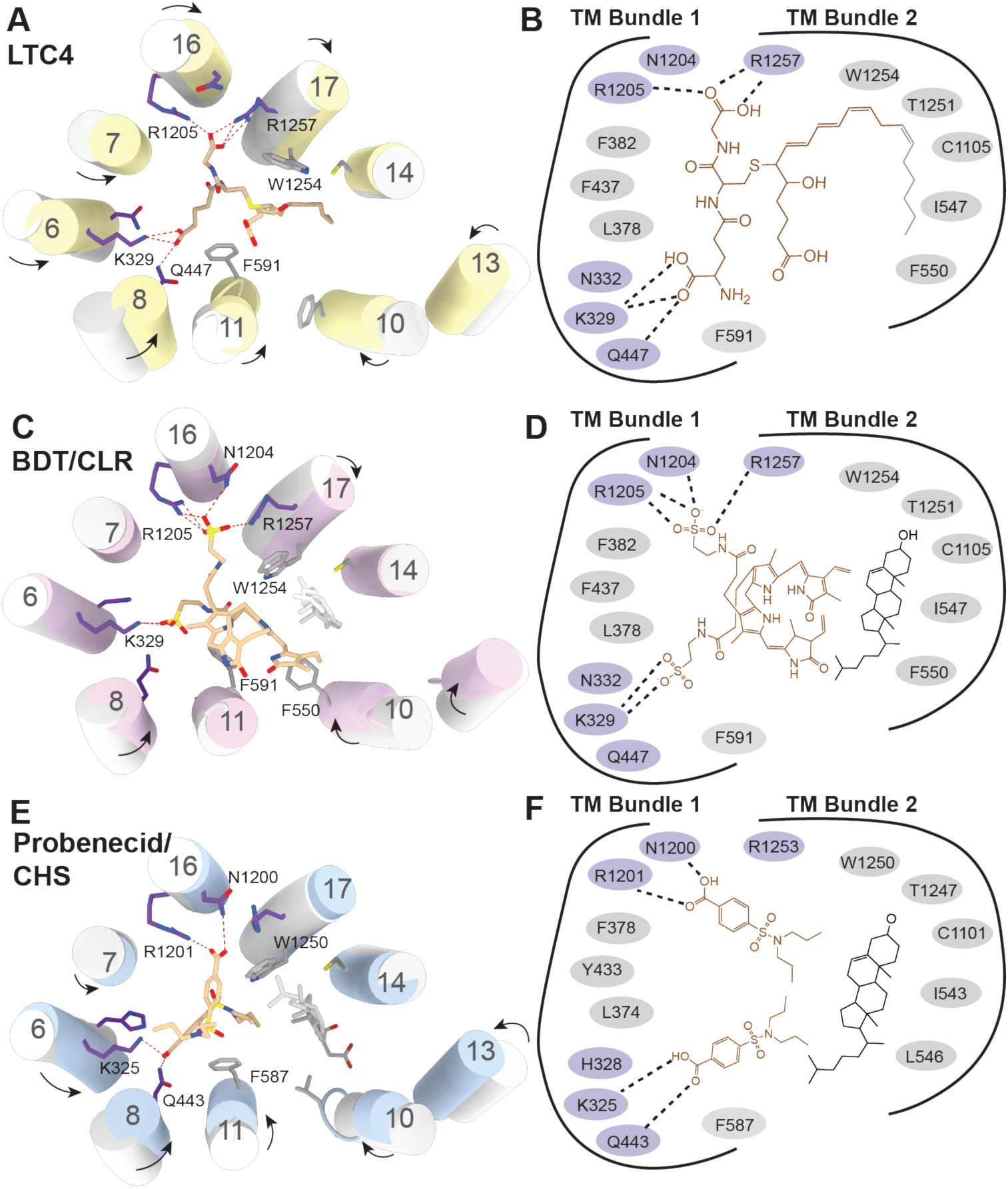
Structural basis of multi-substrate recognition. (A) The structure of the substrate binding site observed in the LTC4-bound conformation (yellow). The structure of the apo form is shown in gray. Residues in the P-pocket that interact with LTC4 are indicated. Hydrogen bonds are depicted as dotted lines. Arrows depict conformational changes upon LTC4 binding. (B) Schematic drawing of the LTC4 binding site. (C) The structure of substrate-binding site observed in the BDT/cholesterol (CLR)-bound conformation (pink). The structure of the apo form is shown in gray. BDT is depicted in tan, CLR in gray. (D) Schematic drawing of the interactions of BDT (tan) and CLR (black) with MRP2. (E) The structure of binding site observed in the Probenecid/cholesterol hemisuccinate (CHS)-bound rMrp2 (blue). The structure of rMrp2 apo form is shown in gray. Probenecid is depicted in tan, CHS in gray. (F) Schematic drawing of the rMrp2 bound with probenecid (tan) and CHS (black). Portion of CHS which extends away from the substrate binding site residues is not pictured.

## Methods

### Cell culture

*Spodoptera frugiperda* (Sf9) cells (Gibco) were cultured in Sf-900 II SFM medium (Gibco) containing 5% fetal bovine serum (FBS) (Gibco) and 1% antibiotic-antimycotic (Gibco) at 27°C. HEK293S GnTl^−^ cells (ATCC) were cultured in Freestyle 293 medium (Gibco) containing 2% FBS and 1% anti-anti at 37°C, 8% CO_2_, and 80% humidity. Cell lines were tested for mycoplasma contamination using the University Mycoplasma Detection Kit (ATCC) every month.

### Protein expression and purification

Mammalian codon-optimized gene encoding human MRP2 (BioBasic Inc.) was subcloned into a plasmid in frame with a C-terminal PreScission Protease cleavage site followed by an eGFP tag. The vector was then transformed into DH10Bac cells to generate recombinant bacmid. Purified bacmid was transfected into Sf9 cells and the resulting baculovirus was amplified to P4. P4 baculovirus was added at 10% v/v to HEK293S GnTl^−^ cells at a density of 2.5-3.5 million cells/mL. Sodium butyrate was added 12 hours later to a final concentration of 10 mM, and the temperature was dropped to from 37 °C to 30 °C. Cells were harvested 48 hours later, flash frozen in liquid nitrogen, and stored at −70 °C.

For protein purification, cells were thawed and solubilized in buffer containing 300 mM NaCl, 50 mM HEPES pH 8.0 with KOH, 2 mM MgCl_2_, 2 mM DTT, 20% glycerol, 2% LMNG, 0.2% CHS, 1 μg/mL aprotinin, 0.6 mM benzamidine, 1 mM PMSF, 1 μg/mL leupeptin, 1 μg/mL pepstatin A, 100 μg/mL soy trypsin inhibitor, and DNase I. Insoluble material was removed by centrifugation at 75,000 xg for 40 minutes. Supernatant was mixed with GFP nanobody-conjugated Sepharose 4 Fast Flow resin (GE Healthcare). Resin was washed with buffer containing 0.06% digitonin, 150 mM NaCl, 50 mM HEPES pH 8.0 with KOH, 2 mM MgCl_2_, 2 mM DTT, and 5% glycerol. PreScission protease was added to the column and incubated overnight at 4°C to release MRP2 from the resin. Eluate was passed through GST Sepharose resin (Cytiva) to remove PreScission protease. The protein was concentrated using a 100 kDa filter to approximately 1 mL and further purified by size-exclusion chromatography at 4°C using a Superose 6 10/300 GL column (GE Healthcare) equilibrated with buffer containing 0.06% digitonin, 150 mM KCl, 50 mM Tris pH 8.0, 2 mM MgCl_2_, and 2 mM DTT.

### Mutagenesis

MRP2(E1462Q) and MRP2(R928A/K932A) were generated using PCR amplification with mutagenic primers. These primers were complementary to the template plasmid except at the bases targeted for mutation (Integrated DNA Technologies, IDT). Following PCR amplification of the MRP2 wild type plasmid with these primers, the product was digested by Dpn1 for 4 hours (New England Biotechnologies, NEB). The PCR product was purified by gel purification (Zymo Research). The purified plasmid was added to 50 uL DH5α cells (NEB) and transformed following the manufacturer protocol. Following recovery, 200 uL transformed cells were spread on LB/ampicillin plates and incubated at 37°C overnight. Clonal plasmid DNA was then expanded, purified, and sequenced for verification (Genewiz).

To generate the Scramble construct, the R-domain amino acid sequence was randomly scrambled and ordered as gene fragments from IDT. The parent MRP2 plasmid was amplified using primers complementary to the regions of MRP2 directly adjacent the R-domain and to the N- and C-termini of the scrambled R-domain. The PCR product of this amplification was purified by gel extraction and assembled with the scrambled R-domain using Gibson assembly (NEB) at a 1:5 g/g and 1:10 g/g ratio. 5 uL of the assembly product was added to 50 uL NEB5a cells, transformed, expanded, and purified as above.

### ATP Hydrolysis assay

ATP hydrolysis activity was measured using the NADH-coupled assay described previously^27,28^. Briefly, MRP2 was added at a concentration of 75 nM to a reaction mixture containing 50 mM HEPES, 150 mM KCl, 2 mM MgCl_2_, 2 mM DTT, 0.06% w/v digitonin, 60 μg/mL pyruvate kinase, 32 μg/mL lactate dehydrogenase, 9 mM phosphoenolpyruvate, and 150 μM NADH in gel filtration buffer. For LTC4 stimulation assays, LTC4 was added to this reaction buffer at no higher than 1.6% DMSO (DMSO volume was consistent across all assayed wells). 30 μL aliquots were distributed into wells of a black/clear bottom Corning 384-well polystyrene microplate. The reaction was initiated by the addition of 4 mM ATP-Mg^2+^ and NADH depletion was measured by monitoring λex/λem 340/445 nm using a Tecan Infinite M1000 Pro (Tecan). The rate of NADH depletion was measured from a minimum of three separate measurements per condition, and converted to ATP turnover with an NADH standard curve. Data was fit to the quadratic velocity equation, a modification of the Michaelis-Menten equation which accounts for tight-binding substrates.

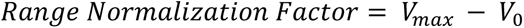

Where *V*_0_ = ATPase activity in absence of LTC4, and *V_max_* = ATPase activity in presence of saturating LTC4.

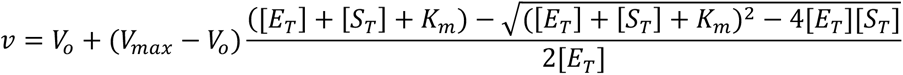

### Cryo-EM sample preparation

MRP2 Apo (PDB:9C21): Immediately following gel filtration, MRP2 was concentrated to 4.5 mg/mL (27 μM). 3 mM fluorinated Fos-choline 8 (FFOS8) was added to sample just before application onto glow-discharged Quantifoil R0.6/1 300 mesh Au grids. 3 uL of sample was applied to grids, blotted for 3 seconds with a blot force of 15 at 100% humidity and 22°C, and frozen in ethane using a Vitrobot Mark IV (Field Electron and Ion Company). The LTC4-bound structure (PDB: 9C21) was obtained in the presence of 285 μM LTC4 and 2.5% DMSO. The MRP2 (E1462Q) structures (9BR2, 9BUK) were obtained in the presence of 5 mM ATP-Mg^2+^ and 40 μM LTC4.

### Data collection

Cryo-EM images for WT MRP2 apo and MRP2(E1462Q) structures were collected as described previously on a Titan Krios (FEI) using a K3 Summit direct electron detector (Gatan, Inc).^20,27^ Cryo-EM images for the LTC4-bound MRP2 were collected on a Titan Krios (FEI) using a Falcon 4i Direct Electron Detector (ThermoFisher). All micrographs were collected in SerialEM using superresolution mode.^29^ Data collection parameters have been summarized in Table S1.

### Image processing

Strategies for data processing have been outlined in supplementary figures 1, 2, and 4. Super-resolution images were corrected for gain reference and binned by 2. Beam-induced motion was corrected using MotionCor2.^30^ Contrast transfer function (CTF) estimation was conducted using CTFFIND4.^31^ Particles were picked using RELION Laplacian-of-Gaussian autopicking function, binned by 4, extracted in RELION and imported into cryoSPARC.^32^ Particles went through several rounds of 2D classification, and resulting particles underwent *ab initio* reconstruction with 3-4 classes. The best class demonstrated density protruding from the detergent micelle, while other classes resembled empty micelles. These classes were used to conduct heterogeneous refinement on the original particle stack. Nonuniform refinement of the best class following heterogeneous refinement yielded a low-resolution reconstruction of MRP2. The particles from the best class were imported into Relion using the csparc2star.py script, where they were re-extracted without binning and sorted by iterations of 3D classification using the density from nonuniform refinement as the reference map.^33^ The best classes were refined by non-uniform refinement and local refinement in cryoSPARC. FSC curves were generated in cryoSPARC. All resolutions are reported at the 0.143 FSC cutoff.

### Model Building and Refinement

The initial models of each conformation of MRP2 were obtained by docking the previously reported models of the transporter core of bMRP1 (PDB:5UJ9, 6BHU) and the AlphaFold2 model of TMD0 into the sharpened maps and mutating each residue to the corresponding MRP2 residue based on sequence alignment.^9,20,34,35^ Models were then manually adjusted into the density using Coot. Models were then refined using ISOLDE and PHENIX, followed by manual adjustment of side chain geometry to best fit the cryo-EM density.^36,37^ Regions with poor side chain density were built as poly alanine models. The quality of the models was evaluated by MolProbity.^38^

## Data availability

The cryo-EM maps of Apo MRP2, inward-facing and outward-facing MRP2(E1462Q), and LTC4-bound MRP2 have been deposited in the Electron Microscopy Data Bank under the accession codes EMD-45159, EMD-44833, EMD-44911, and EMD-45099, respectively. The corresponding atomic models have been deposited in the Protein Data Bank under accession codes 9C2I, 9BR2, 9BUK, and 9C12. All other data are available in the main text or in supplemental information.

## Acknowledgments

We thank Mark Ebrahim, Johanna Sotiris, and Honkit Ng at the Rockefeller University Evelyn Gruss Lipper Cryo-EM Resource Center and Nick Spellmon at the HHMI Janelia Cryo-EM Facility for their support with collecting electron microscopy data. We thank members of the Chen and Mackinnon laboratories for helpful discussions. This work was supported by HHMI to J.C., the National Institute of General Medical Sciences (T32GM007739 to the Weill Cornell/Rockefeller/Sloan Kettering Tri-Institutional MD-PhD program) and a Predoctoral Fellowship from the National Cancer Institute (F30CA257282) to H.L.P.

## Author Contributions

E.K. and J.C. designed research; E.K. performed all the experiments with help from H.L.P., and J.B.; E.K. and J.C. analyzed data; E.K. and J.C. wrote the paper.

## References

1. Cole, S. P. C. & Deeley, R. G. Multidrug resistance mediated by the ATP-binding cassette transporter protein MRP. BioEssays 20, 931–940 (1998).

2. Gerk, P. M. & Vore, M. Regulation of Expression of the Multidrug Resistance-Associated Protein 2 (MRP2) and Its Role in Drug Disposition. J. Pharmacol. Exp. Ther. 302, 407–415 (2002).

3. Paulusma, C. C. et al. A mutation in the human canalicular multispecific organic anion transporter gene causes the Dubin-Johnson syndrome. Hepatology 25, 1539–1542 (1997).

4. Keitel, V. et al. A common Dubin-Johnson syndrome mutation impairs protein maturation and transport activity of MRP2 (ABCC2). Am. J. Physiol.-Gastrointest. Liver Physiol. 284, G165–G174 (2003).

5. Lee, J.-H. et al. Neonatal Dubin-Johnson Syndrome: Long-Term Follow-up and MRP2 Mutations Study. Pediatr. Res. 59, 584–589 (2006).

6. Choi, J. H. et al. MRP2 haplotypes confer differential susceptibility to toxic liver injury. Pharmacogenet. Genomics 17, 403–415 (2007).

7. Vlaming, M. L. H. et al. Carcinogen and Anticancer Drug Transport by Mrp2 in Vivo: Studies Using Mrp2 (Abcc2) Knockout Mice. J. Pharmacol. Exp. Ther. 318, 319–327 (2006).

8. Yamasaki, M. et al. Role of multidrug resistance protein 2 (MRP2) in chemoresistance and clinical outcome in oesophageal squamous cell carcinoma. Br. J. Cancer 104, 707–713 (2011).

9. Johnson, Z. L. & Chen, J. Structural Basis of Substrate Recognition by the Multidrug Resistance Protein MRP1. Cell 168, 1075–1085.e9 (2017).

10. Bakos, É. et al. Functional Multidrug Resistance Protein (MRP1) Lacking the N-terminal Transmembrane Domain*. J. Biol. Chem. 273, 32167–32175 (1998).

11. Alam, A. & Locher, K. P. Structure and Mechanism of Human ABC Transporters. Annu. Rev. Biophys. 52, 275–300 (2023).

12. Alam, A., Kowal, J., Broude, E., Roninson, I. & Locher, K. P. Structural insight into substrate and inhibitor discrimination by human P-glycoprotein. Science 363, 753–756 (2019).

13. Nosol, K. et al. Cryo-EM structures reveal distinct mechanisms of inhibition of the human multidrug transporter ABCB1. Proc. Natl. Acad. Sci. 117, 26245–26253 (2020).

14. Manolaridis, I. et al. Cryo-EM structures of a human ABCG2 mutant trapped in ATP-bound and substrate-bound states. Nature 563, 426–430 (2018).

15. Jackson, S. M. et al. Structural basis of small-molecule inhibition of human multidrug transporter ABCG2. Nat. Struct. Mol. Biol. 25, 333–340 (2018).

16. Orlando, B. J. & Liao, M. ABCG2 transports anticancer drugs via a closed-to-open switch. Nat. Commun. 11, 2264 (2020).

17. Kowal, J. et al. Structural Basis of Drug Recognition by the Multidrug Transporter ABCG2. J. Mol. Biol. 433, 166980 (2021).

18. Oldham, M. L. et al. A mechanism of viral immune evasion revealed by cryo-EM analysis of the TAP transporter. Nature 529, 537–540 (2016).

19. Oldham, M. L., Grigorieff, N. & Chen, J. Structure of the transporter associated with antigen processing trapped by herpes simplex virus. eLife 5, e21829 (2016).

20. Johnson, Z. L. & Chen, J. ATP Binding Enables Substrate Release from Multidrug Resistance Protein 1. Cell 172, 81–89.e10 (2018).

21. Wang, L. et al. Characterization of the kinetic cycle of an ABC transporter by single-molecule and cryo-EM analyses. eLife 9, e56451.

22. Cui, Y. et al. Drug Resistance and ATP-Dependent Conjugate Transport Mediated by the Apical Multidrug Resistance Protein, MRP2, Permanently Expressed in Human and Canine Cells. Mol. Pharmacol. 55, 929–937 (1999).

23. Grant, C. E., Gao, M., DeGorter, M. K., Cole, S. P. C. & Deeley, R. G. Structural Determinants of Substrate Specificity Differences between Human Multidrug Resistance Protein (MRP) 1 (ABCC1) and MRP3 (ABCC3). Drug Metab. Dispos. 36, 2571–2581 (2008).

24. Mao, Y.-X. et al. Transport mechanism of human bilirubin transporter ABCC2 tuned by the inter-module regulatory domain. Nat. Commun. 15, 1061 (2024).

25. Mazza, T. et al. Structural basis for the modulation of MRP2 activity by phosphorylation and drugs. Nat. Commun. 15, 1983 (2024).

26. Bakos, É. et al. Interactions of the Human Multidrug Resistance Proteins MRP1 and MRP2 with Organic Anions. Mol. Pharmacol. 57, 760–768 (2000).

27. Pietz, H. L. et al. A macrocyclic peptide inhibitor traps MRP1 in a catalytically incompetent conformation. Proc. Natl. Acad. Sci. 120, e2220012120 (2023).

28. Scharschmidt, B. F., Keeffe, E. B., Blankenship, N. M. & Ockner, R. K. Validation of a recording spectrophotometric method for measurement of membrane-associated Mg- and NaK-ATPase activity. J. Lab. Clin. Med. 93, 790–799 (1979).

29. Mastronarde, D. N. Automated electron microscope tomography using robust prediction of specimen movements. J. Struct. Biol. 152, 36–51 (2005).

30. Zheng, S. Q. et al. MotionCor2: anisotropic correction of beam-induced motion for improved cryo-electron microscopy. Nat. Methods 14, 331–332 (2017).

31. Rohou, A. & Grigorieff, N. CTFFIND4: Fast and accurate defocus estimation from electron micrographs. J. Struct. Biol. 192, 216–221 (2015).

32. Zivanov, J. et al. New tools for automated high-resolution cryo-EM structure determination in RELION-3. eLife 7, e42166 (2018).

33. Asarnow, D., Palovcak, E., Cheng, Y. UCSF pyem v0.5. Zenodo 10.5281/zenodo.3576630 (2019).

34. Jumper, J. et al. Highly accurate protein structure prediction with AlphaFold. Nature 596, 583–589 (2021).

35. Varadi, M. et al. AlphaFold Protein Structure Database in 2024: providing structure coverage for over 214 million protein sequences. Nucleic Acids Res. 52, D368–D375 (2024).

36. Croll, T. I. ISOLDE: a physically realistic environment for model building into low-resolution electron-density maps. Acta Crystallogr. Sect. Struct. Biol. 74, 519–530 (2018).

37. Liebschner, D. et al. Macromolecular structure determination using X-rays, neutrons and electrons: recent developments in Phenix. Acta Crystallogr. Sect. Struct. Biol. 75, 861–877 (2019).

38. Chen, V. B. et al. MolProbity: all-atom structure validation for macromolecular crystallography. Acta Crystallogr. D Biol. Crystallogr. 66, 12–21 (2010).

