## Supplemental files for "Structural basis for the transport and regulation mechanism of the Multidrug resistance-associated protein 2"

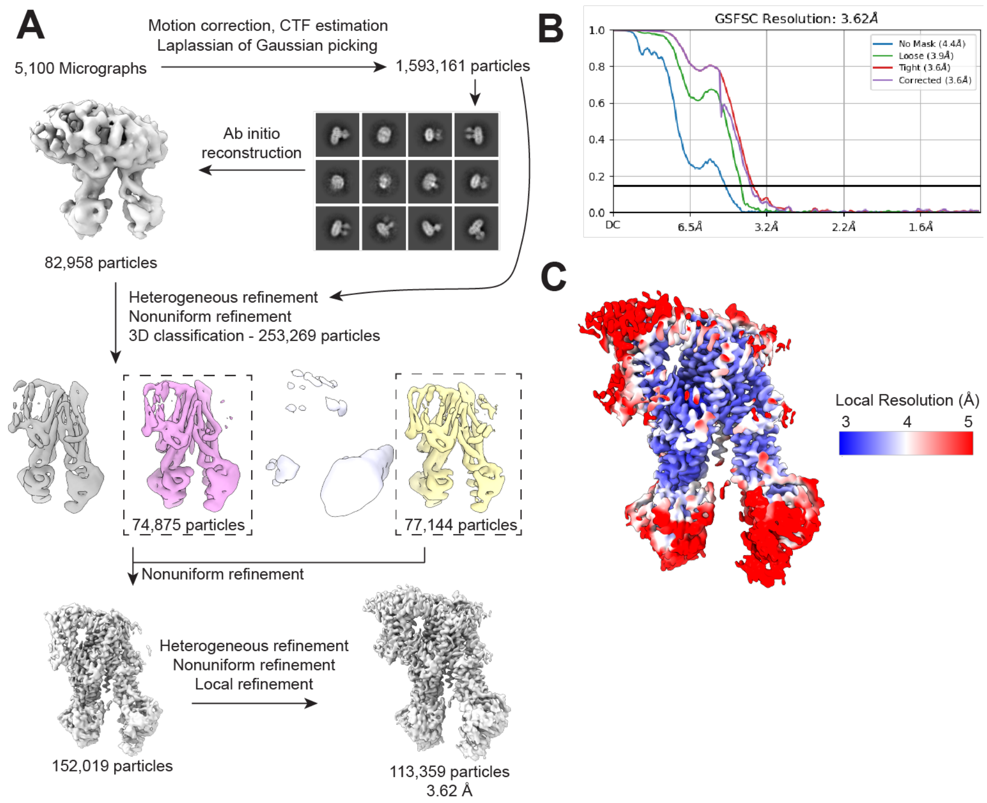

**Supplementary Figure 1. Structure determination of the ligand-free MRP2.**

(A) The image processing workflow.

(B) Fourier shell correlation (FSC) curves of the final map generated in CryoSPARC. Blue: no mask applied; green: with a soft solvent mask: red: with a tight mask around the protein; and purple: “Corrected” represents FSC calculated after applying the tight mask and correcting by noise substitution (Chen et al 2013 Ultramicroscopy).

(C) Local resolution estimation of the final reconstruction.

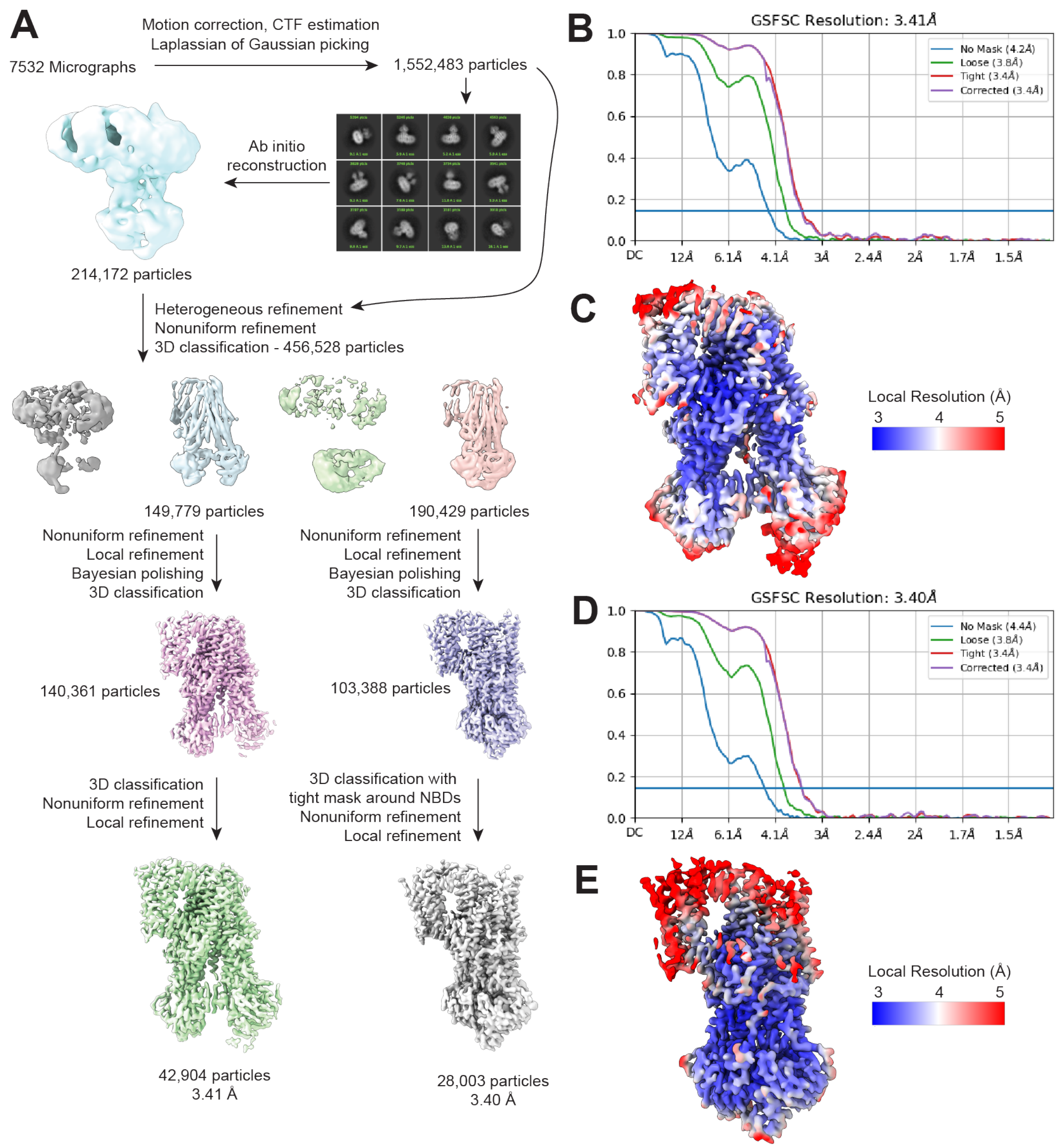

**Supplementary Figure 2. Structure determination of the E1462Q variant in the presence of ATP.**
(A) Image processing workflow.
(B) Fourier shell correlation curves of the inward-facing structure.
(C) Local resolution estimation for the inward-facing structure.
(D) Fourier shell correlation curves of the NBD-dimerized conformation.
(E) Local resolution estimation for the NBD-dimerized conformation.

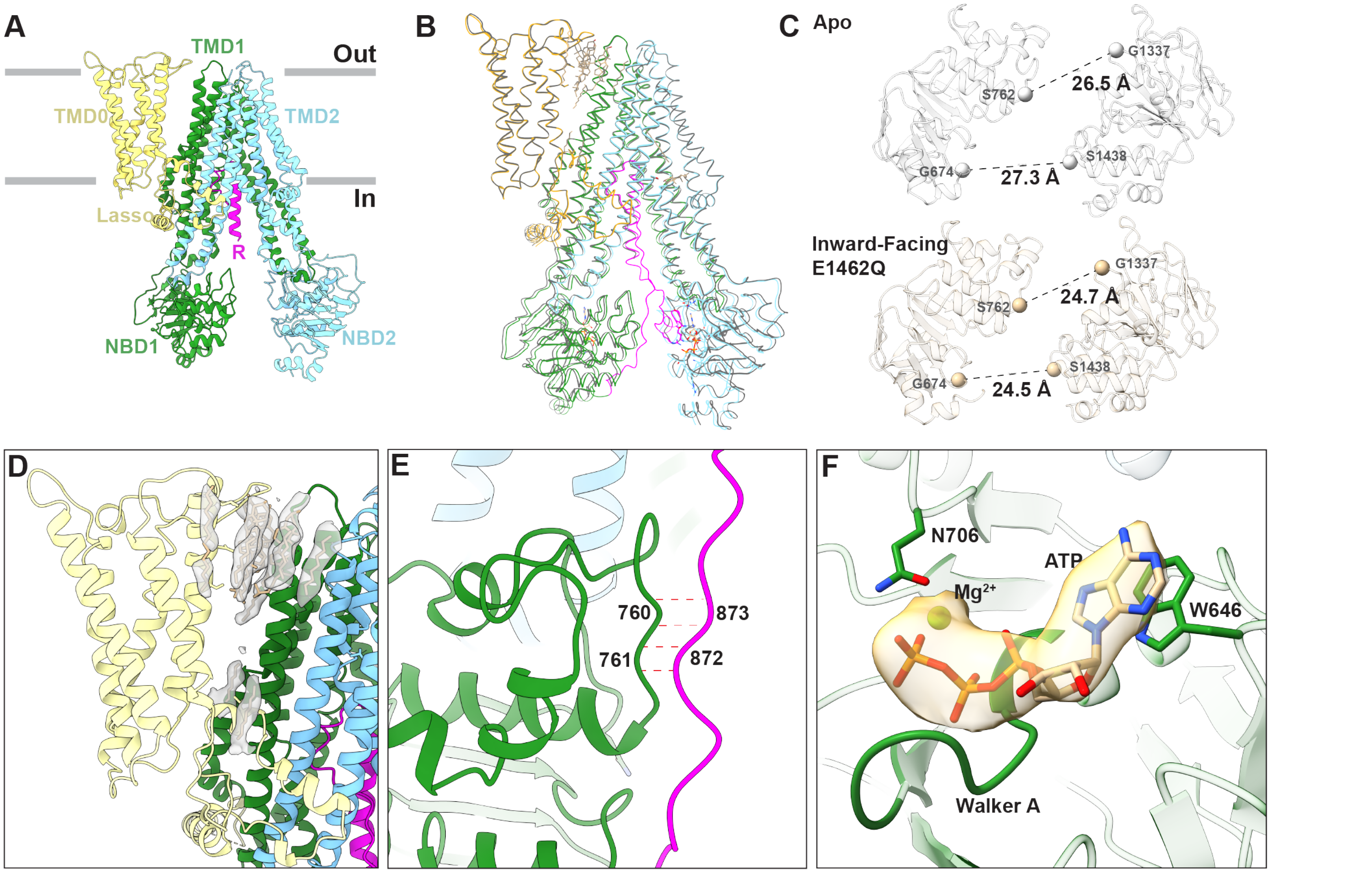

**Supplemental Figure 3. Two similar auto-inhibited structures**

(A) The overall structure of the WT MRP2 in the ligand-free, apo conformation.
(B) Structural superposition of the E1462Q inward-facing conformation (colored) and the WT apo conformation (gray).
(C) Structural comparison of the NBDs of observed in the WT apo and E1462Q inward-facing conformations. Distances between pairs of ATP binding residues are indicated.
(D) Structure of the TMD0 observed in the MRP2(E1462Q) reconstruction, with cryo-EM density corresponding to cholesterol and lipid depicted as surface.
(E) Residues 758-762 of NBD1 closely interact with the R domain as it extends from NBD1, forming a beta sheet with residues 760 and 761.
(F) A unique ATP-binding site observed in the MRP2(E1462Q) structure. Cryo-EM density for ATP and magnesium are shown as yellow surface.

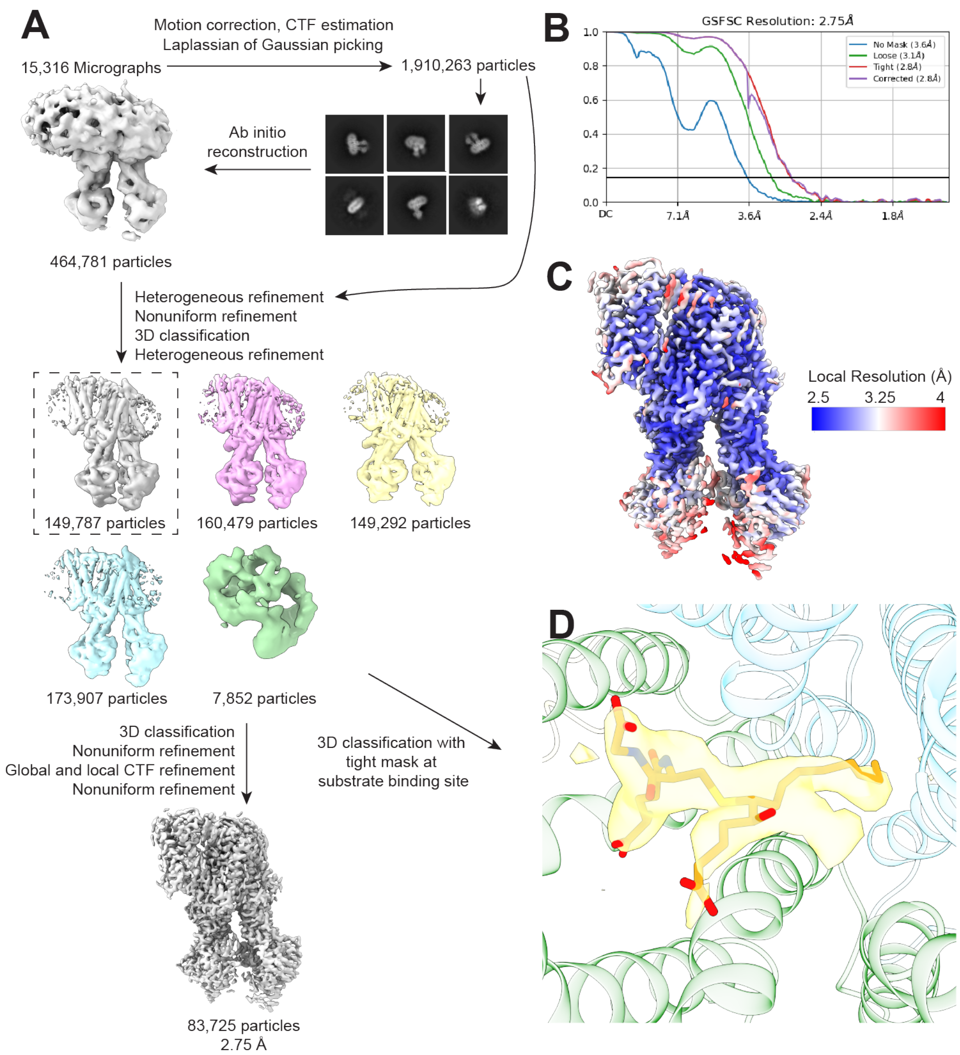

**Supplementary Figure 4. Structure determination of the LTC4-bound structure.**

(A) Image processing workflow.
(B) Fourier shell correlation curves.
(C) Local resolution estimation.
(D) Density (yellow) corresponding to LTC4 at the substrate-binding site. The hydrophobic tail of LTC4 is removed as no clear density was observed in this region.

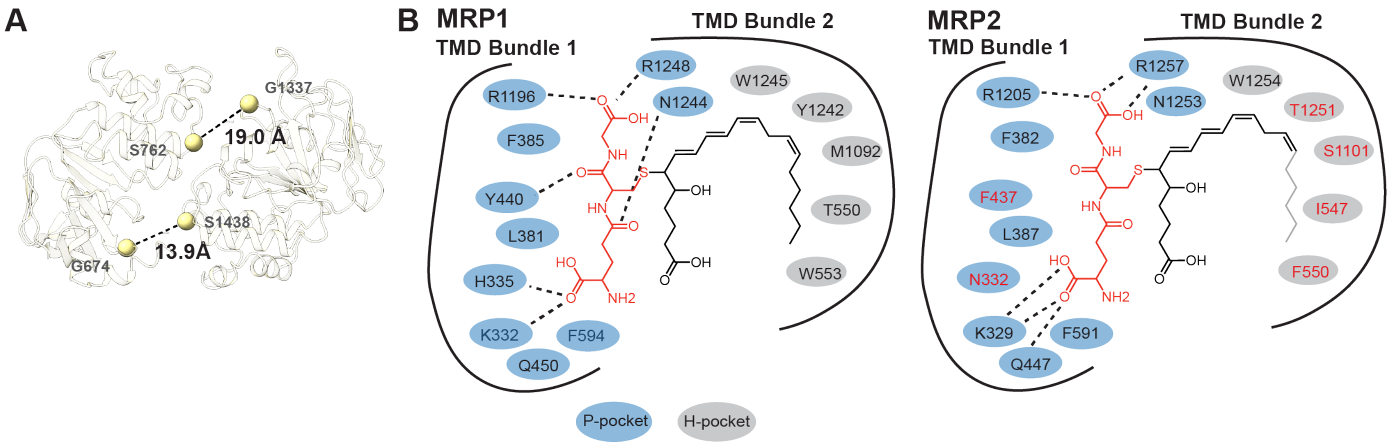

**Supplemental figure 5. The LTC4-bound conformation**

(A) The structure of the NBDs observed in the LTC4-bound MRP2 reconstruction.
(B) Schematic drawings comparing LTC4-binding in MRP1 and MRP2**.** The polar P-pocket residues of MRP1 and their corresponding residues in MRP2 are depicted in blue, and hydrophobic H-pocket residues of MRP1 and their corresponding residues in MRP2 are depicted in gray. Conserved residues are written in black, differing residues are written in red. Black dotted lines indicate hydrogen bonding. Unresolved hydrophobic tail of LTC4 is depicted in light gray.

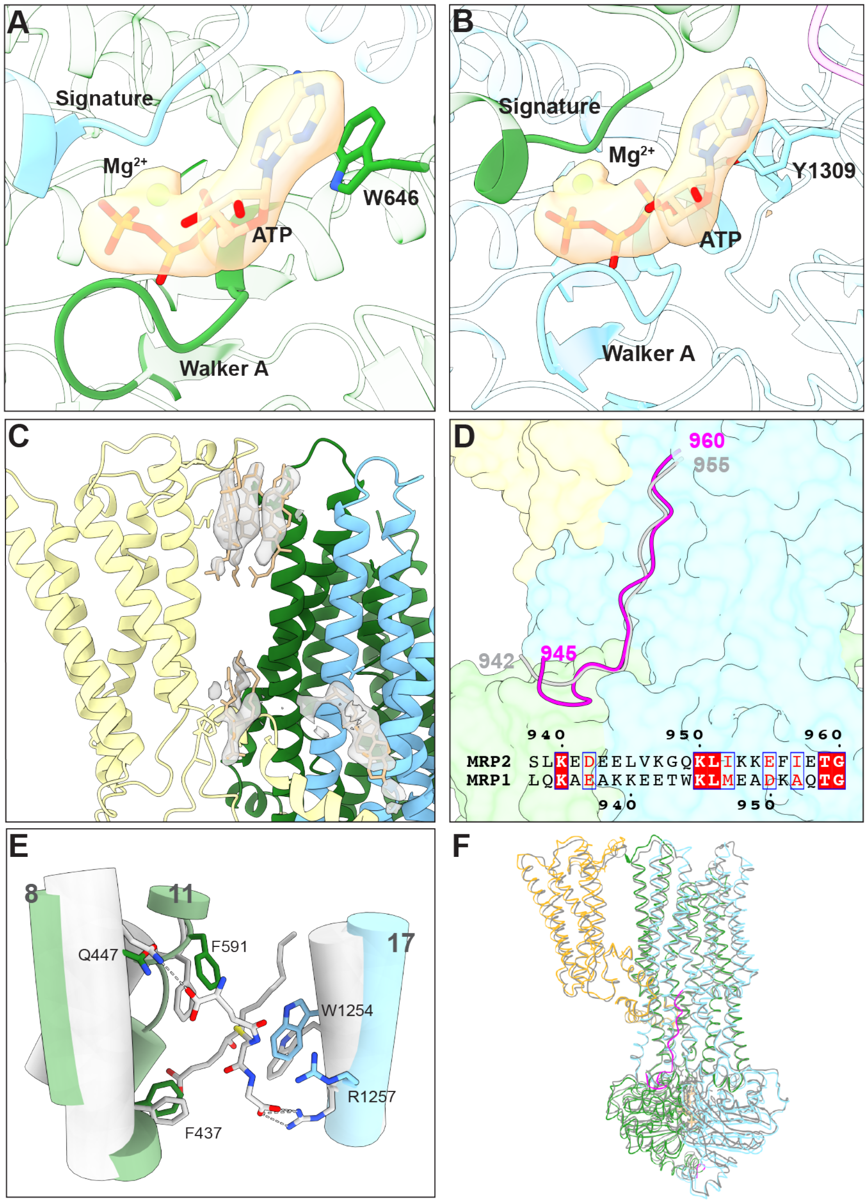

**Supplementary figure 6. The structure of MRP2(E1462Q) in the ATP-bound, NBD-dimerized conformation**

(A) Structure of the degenerate ATPase site. Residues from NBD1 are depicted in green, residues from NBD2 are depicted in blue. Cryo-EM density of ATP and magnesium are depicted as tan surface.
(B) Structure of the consensus ATPase site.
(C) Zoomed-in view of the TMD0 region, with cryo-EM density corresponding to cholesterol and lipids depicted as surface.
(D) Structural comparison of the MRP2 R domain (magenta) and MRP1 linker domain (gray), along with amino acid sequence alignment.
(E) Conformational changes at the substrate-binding site. The LTC4-bound structure is depicted in gray and the outward-facing structure depicted in green (TMD1) and blue (TMD2).
(F) Structural superposition of the MRP2(E1462Q) ATP-bound, pre-hydrolysis conformation (colored) with that of the WT MRP2 in the ATP/ADP-bound, post-hydrolysis conformation (PDB: 8JXU)

**Supplementary Table 1. Data collection, processing, and structure refinement**

| | MRP2(E1462Q) inward-facing (EMDB-44833) (PDB 9BR2) | MRP2(E1462) Outward-facing (EMDB-44911) (PDB 9BUK) | MRP2 285 μM LTC4 (EMDB-45099) (PDB 9C12) | MRP2 Apo (EMDB-45159) (PDB 9C2I) |
|---|---|---|---|---|
| **Data collection and processing** | | | | |
| Magnification | 105,000 | 105,000 | 165,000 | 105,000 |
| Voltage (kV) | 300 | 300 | 300 | 300 |
| Electron exposure (e–/Å$^2$) | 65 | 65 | 65 | 65 |
| Defocus range (μm) | 0.8 to 2.5 | 0.8 to 2.5 | 0.8 to 1.6 | 0.8 to 2.5 |
| Pixel size (Å) | 0.676 | 0.676 | 0.743 | 0.676 |
| Symmetry imposed | C1 | C1 | C1 | C1 |
| Initial particle images (no.) | 1,552,483 | 1,552,483 | 1,910,263 | 1,593,161 |
| Final particle images (no.) | 42,904 | 28,003 | 83,725 | 113,359 |
| Map resolution (Å) | 3.41 | 3.40 | 2.75 | 3.62 |
| FSC threshold | 0.143 | 0.143 | 0.143 | 0.143 |
| **Refinement** | | | | |
| Model resolution (Å) | 3.4 | 3.4 | 2.9 | 3.6 |
| FSC threshold | 0.143 | 0.143 | 0.143 | 0.143 |
| Model resolution range (Å) | 3.2 to 3.8 | 4.3 to 3.7 | 2.7 to 3.7 | 3.5 to 4.0 |
| Map sharpening *B* factor (Å$^2$) | 70.7 | 62.7 | 58.4 | 116.4 |
| Model composition | | | | |
| Non-hydrogen atoms | 11493 | 10815 | 10983 | 11186 |
| Protein residues | 1469 | 1394 | 1398 | 1422 |
| Ligands | Magnesium: 1<br>ATP: 2<br>CLR: 5<br>UNL: 8 | Magnesium: 2<br>ATP: 2<br>CLR: 5<br>UNL: 4 | LTC4: 1<br>CLR: 5<br>UNL: 13 | CLR: 5<br>UNL: 8 |
| *B* factors (Å$^2$) | | | | |
| Protein | 66.40 | 56.86 | 104.86 | 65.58 |
| Ligand | 102.58 | 86.01 | 95.26 | 59.61 |
| R.m.s. deviations | | | | |
| Bond lengths (Å) | 0.009 (0) | 0.007 (1) | 0.012 (4) | 0.009 (0) |
| Bond angles (°) | 1.083 (18) | 0.937 (10) | 1.745 (59) | 1.060 (13) |
| Validation | | | | |
| MolProbity score | 1.53 | 1.41 | 0.99 | 1.57 |
| Clashscore | 3.78 | 3.15 | 2.16 | 4.24 |
| Poor rotamers (%) | 0 | 0 | 0 | 0 |
| Ramachandran plot | | | | |
| Favored (%) | 94.67% | 95.59% | 98.13% | 94.77% |
| Allowed (%) | 5.33% | 4.41% | 1.87% | 5.23% |
| Disallowed (%) | 0 | 0 | 0 | 0 |
